# Uncovering of cytochrome P450 anatomy by SecStrAnnotator

**DOI:** 10.1101/2020.04.15.042531

**Authors:** Adam Midlik, Veronika Navrátilová, Taraka Ramji Moturu, Jaroslav Koča, Radka Svobodová, Karel Berka

**Affiliations:** CEITEC – Central European Institute of Technology, Masaryk University, Brno 625 00, Czech Republic; National Centre for Biomolecular Research, Faculty of Science, Masaryk University, Brno 625 00, Czech Republic; Department of Physical Chemistry, Faculty of Science, Palacký University, Olomouc 771 46, Czech Republic

## Abstract

Protein structural families are groups of homologous proteins defined by the organization of secondary structure elements (SSEs). Nowadays, many families contain vast numbers of structures, and the SSEs can help to orient within them. Communities around specific protein families have even developed specialized SSE annotations, always assigning the same name to the equivalent SSEs in homologous proteins. A detailed analysis of the groups of equivalent SSEs provides an overview of the studied family and enriches the analysis of any particular protein at hand.

We developed a workflow for the analysis of the secondary structure anatomy of a protein family. We applied this analysis to the model family of cytochromes P450 (CYPs) – a family of important biotransformation enzymes with a community-wide used SSE annotation. We report the occurrence, typical length and amino acid sequence for the equivalent SSE groups, the conservation/variability of these properties and relationship to the substrate recognition sites. We also suggest a generic residue numbering scheme for the CYP family. Comparing the bacterial and eukaryotic part of the family highlights the significant differences and reveals an anomalous group of bacterial CYPs with some typically eukaryotic features. Application of our workflow to other families could produce equally surprising findings.

## Introduction

Secondary structure elements (SSEs) are defined by the repetitive pattern of hydrogen bonds and geometric arrangement. The most well-known SSE types are the α-helix and the β-strand. SSEs have been used to analyze protein structures since their first observation by Linus Pauling^1,2^. They define the structural folds of individual protein structural families as classified by CATH^3^ or SCOPe^4^ databases. Structural folds, defined by SSEs, may reveal a possible biochemical function of proteins from those protein families^5^. SSEs can also serve as guides for orientation in the protein structures within scientific communities.

In order to compare similar structures, the communities around several proteins families have developed specialized nomenclatures for annotation (labeling) of the SSEs in the members of the family. Suc h nomenclatures assign the same label to the equivalent SSEs from different proteins in the family. We will refer to a group of equivalent SSEs as an *SSE class*.

Adoption of such SSE nomenclatures is typical for well-studied protein families with a large number of available structures with a common fold but large sequence variations, such as esterases^6,7^, G-protein coupled receptors (GPCRs)^8^, immunoglobulins^9^, cytochromes P450 (CYPs)^10^ and others. These traditional nomenclatures prove to be particularly useful when comparing existing structures, describing new ones or generalizing observations over the whole family.

Annotated SSEs can be used as reference points to describe the position of key regions, such as catalytic sites, selectivity filters, channels, or protein-protein interfaces. A nice illustrative example is again the CYP family (Fig. 1), with a well-established classification of multiple different channels based on their position relative to the annotated SSEs^11–13^ and with substrate selectivity defining residues on several SSEs^14–18^. The channels of the cytochromes P450 represent a network of the ins and outs which provide the ways for the metabolites to enter the deeply buried active site with the heme cofactor as well as the exit paths for the metabolite output. Channels are named according to their spatial location in respect to SSEs lining each pathway and are summarized in the general nomenclature for cytochrome P450 channels introduced by Cojocaru et al .^11^. These SSEs relate to the regions important for substrate recognition – SRS-1, SRS-2, SRS-3, and SRS-5^17^. Moreover, amino acids located in particular SSEs (e.g. F-G loop) play a crucial role in the substrate egress through the channels. In summary, the spatial arrangement of SSEs may differ from one cytochrome P450 to another, which may result in variations of channel opening. Annotation of the SSEs in various structures of cytochrome P450 is useful for identifying these channels and thus elucidating the channel preferences of individual cytochromes P450 and its substrates.

**Figure 1.**
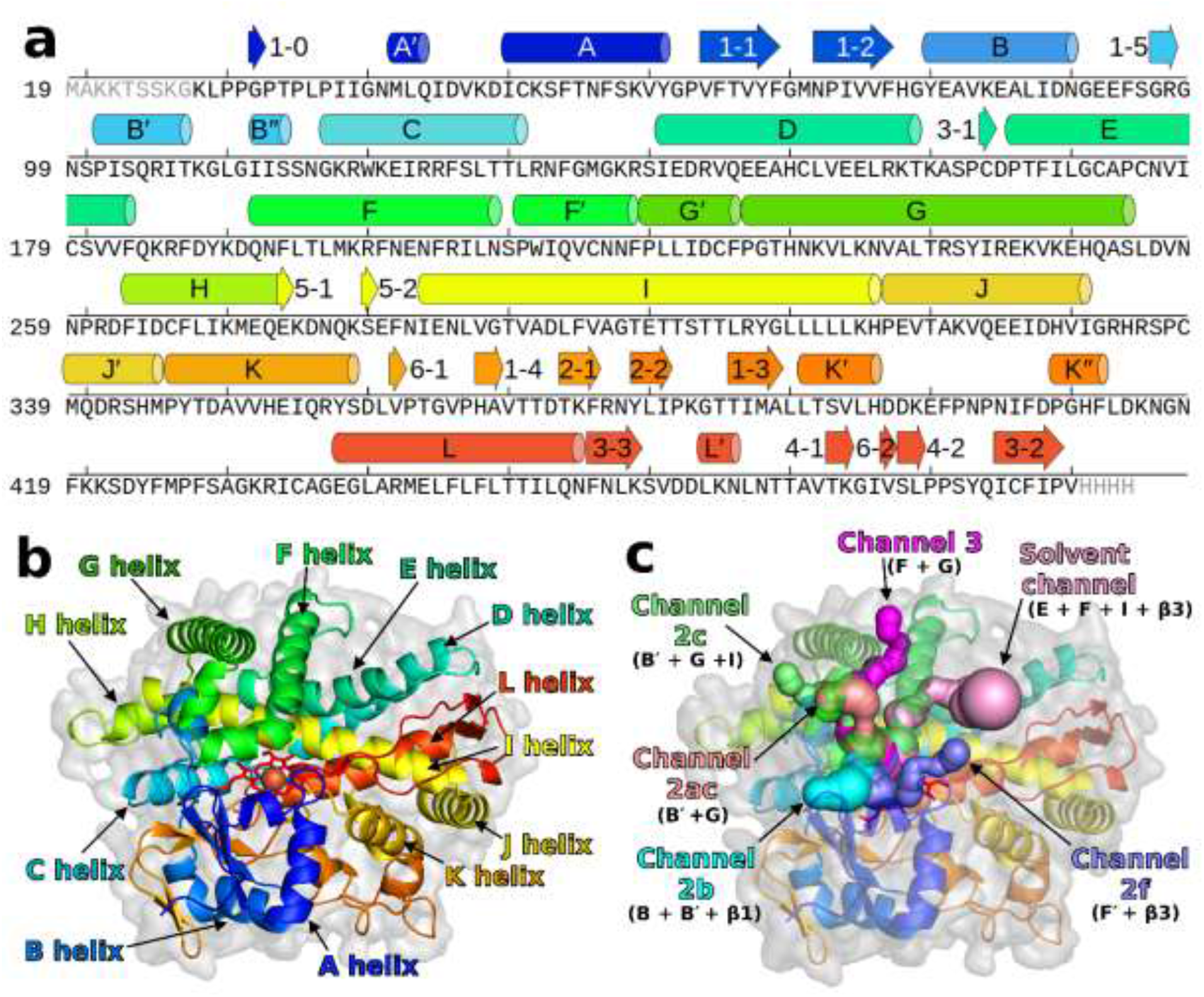
SSE annotation for the CYP family. (**a**) Annotation shown on the sequence (PDB ID 2nnj), (**b**) annotation of the major helices shown on the structure, (**c**) annotation of protein channels defined with respect to the annotated SSEs. Some SSEs (B″, β1-0, β5-1, β5-2, β6-1, β6-2) are not really present in 2nnj but they are shown to illustrate their location in other CYPs. The table of residue ranges of the SSEs is included in Supplementary Table S1.

In some protein families, the communities have extended the annotation down to the residue level and introduced generic numbering schemes. Such schemes assign the same generic number to the equivalent residue positions in homologous proteins, which facilitates comparisons of mutation effects, ligand interactions, structural motifs etc. The generic numberings may be based on the sequence information (e.g. immunoglobulins^19^) or may combine the SSE annotations with the sequence information (e.g. GPCRs^8^).

Annotation of SSEs can be valuable even in protein families without defined traditional nomenclature since it provides the equivalence between the SSEs from the individual members of the family (the SSEs from the same SSE class are annotated by the same label, even if the labels are arbitrarily created).

Previously, we presented methods for automated annotation of SSEs in protein families, implemented in tool SecStrAnnotator^20^. Automated annotation opens the possibility to focus on any protein family and describe its general SSE anatomy, which can bring a valuable insight to the understanding of its function – SSEs typically present, their occurrence, typical length, position and amino acid composition and variation for individual SSE classes.

In this paper, we propose a procedure for analysis of the general SSE anatomy of a protein structural family, based on and extending SecStrAnnotator. We demonstrate this type of analysis on the cytochromes P450, a biologically important family with a long tradition of SSE annotation and well-established SSE nomenclature. We also suggest a generic residue numbering scheme for the CYP family. The SSE annotations for the CYP family are accessible online through SecStrAPI and can be easily visualized via a dedicated PyMOL plugin – all freely available at our website^21^.

### Traditional SSE nomenclature in the CYP family

The common fold of the CYP family has a triangular prism shape and consists mostly of α-helices combined with several β-sheets and a heme cofactor, which forms the catalytic center of the enzyme^14,22,23^. The traditional SSE nomenclature in the CYP family is based on the labels used by Poulos *et al*.^24^ for the first experimentally determined CYP structure (P450cam) with 12 helices, labeled A –L, and 5 sheets, β1–β5. The publication of the refined structure^25^ mentioned a new helix B′ and also several shorter helices and strands without labels.

Ravichandran *et al*.^26^ on P450 BM3 changed the labeling scheme for β-sheets to the form which later became widely used in the community: sheets β1 (5 strands, previously labeled β1+β3), β2 (2 strands, previously β4), β3 (3 strands, previously β5) and a new sheet β4 (2 strands). The strands within each sheet can be referred to individually, using a hyphen (e.g. β1-1, β1-2). They also added annotation of two new helices – J′ (between J and K) and K′ (after strand β1-3).

As more and more structures emerged in the following years, new labels were needed for the newly observed SSE classes: helices A ′^27^, L′^28,29^, F′, G′^30^, K″^29^ and B″^31^; sheets β5^27^ (corresponding to β2 in Poulos *et al*.^25^) and β6^32^; and strand β1-0^33^. Many other SSEs have been mentioned and labeled in literature but these are either very rare or can be treated as a part of a longer SSE (e.g. helix D′ in Park *et al*.^28^ can be understood as an N-terminal part of helix D).

Unfortunately, the labeling is not always consistent, sometimes even in the papers by the same author. The same SSE class can be assigned different labels (e.g. helix L ′ in Scott *et al*.^29^ is helix M in Pylypenko *et al*.^33^) or one label can be assigned to different SSE classes (e.g. helix A ″ is located between β1-1 and β1-2 in Pylypenko *et al*.^33^ but before A′ in Williams *et al*.^34^). Throughout this paper we will use the nomenclature as shown in Fig. 1 and specified in Supplementary Table S1.

## Results and discussion

Structures of proteins from the CYP family typically contain at least 14 helices – A, B, B′, C, D, E, F, G, H, I, J, K, K′, L – and 4 sheets – β1 (5 strands), β2 (2 strands), β3 (3 strands), β4 (2 strands). Additional helices A′, B″, F′, G′, J′, K″ and L′ are often present, as well as two sheets β5 and β6 (with two strands in each). Sheet β1 often contains an extra strand β1-0 (bonded to β1-1). The sequential order and 3D position of these SSE classes is shown in Fig. 1. Other SSEs may appear in the structures, but they are not characteristic to the family.

For convenience, we divide all annotated SSE classes into three groups throughout the paper: *major helices* (helices A–L, typically longer than 8 residues), *minor helices* (all the remaining helices, typically shorter than 8 residues) and *strands*.

In the following text, we analyze these SSE classes in terms of frequency of occurrence, length (number of residues) and amino acid composition. We also discuss differences between eukaryotic and bacterial structures. In Supplementary Note we provide a dedicated analysis of the structural irregularities.

### Regions of variable secondary structure

There are several regions in CYP structures which are structurally very variable. These regions are mainly:

- Region before helix A (containing β1-0 and A′) – this region usually contains at least one helix (annotated as A′), but often there are more short helices. Furthermore, in eukaryotic CYPs this region should contain membrane anchor, but it is missing in most experimental structures, because it complicates crystallization. Therefore, the structures might be biologically irrelevant in this region (though helix A′ was observed also in molecular dynamics simulations on membranes^35^).
- Region between β1-5 and B″ (containing helix B′), is a part of so-called BC-loop. Usually it contains one helix, which is annotated as B′, but often there is more than one helix (especially in bacteria).
- Region between K′ and L (containing helix K″) – this region can contain several short helices with variable positions.

Each of these regions can contain more than one helix and the position of those helices varies from structure to structure. As a result, there is large uncertainty in the annotation of these regions. Therefore, the results for β1-0, A′, B′, K″ should be interpreted with caution.

### Frequency of occurrence of the SSE classes

This section describes the frequency of occurrence of each SSE class, i.e. in what fraction of the structures the particular SSE is present. The results are shown in Fig. 2.

**Figure 2.**
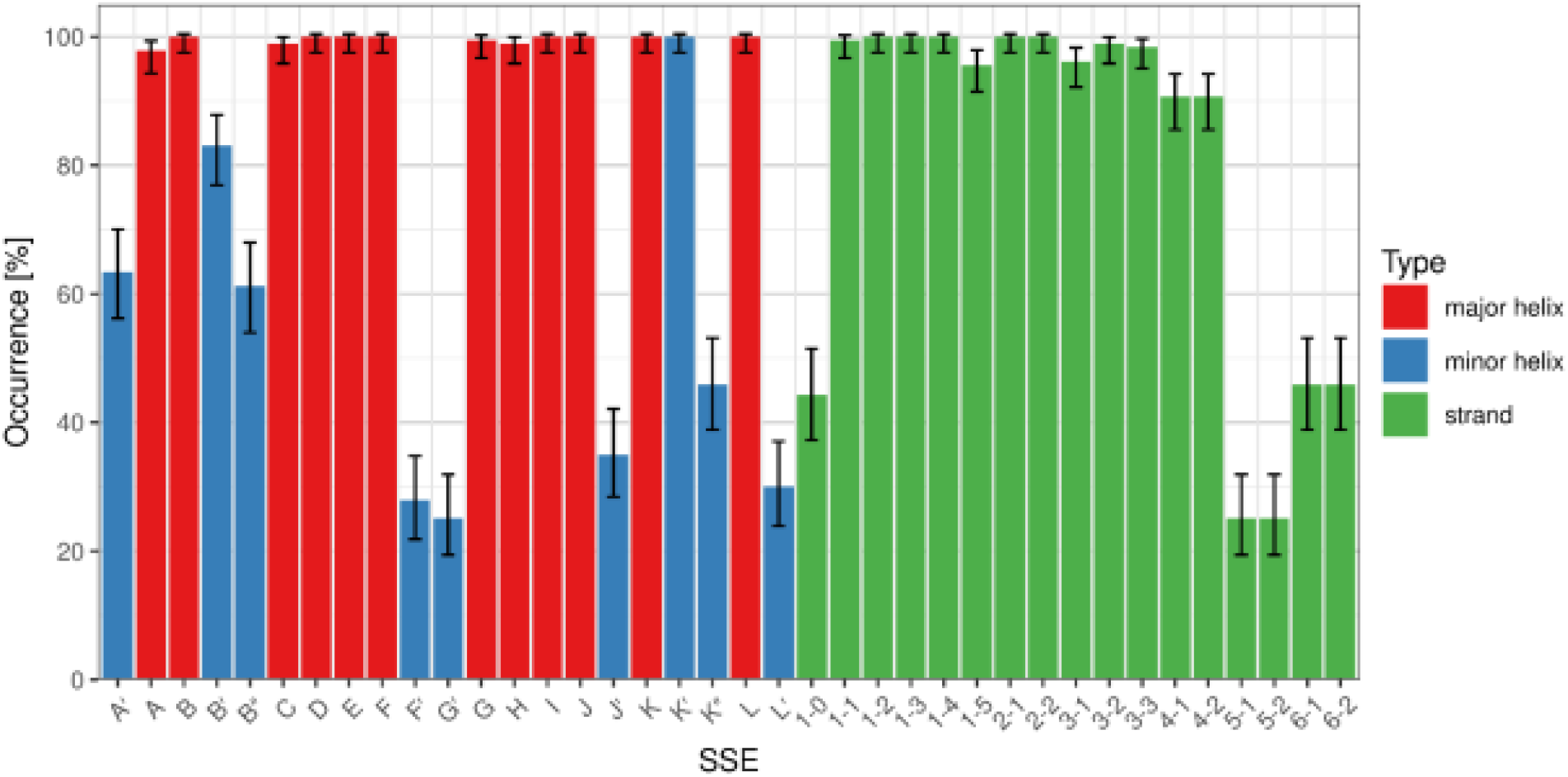
Frequency of occurrence of individual SSE classes. Error bars show confidence intervals calculated by the Agresti-Coull method for α = 0.05^51^.

#### Major helices

All major helices occur in more than 97% of the structures. In cases where they are missing, it can be attributed to the experiment (bad quality, residue coverage or resolution of the structure), rather than not being formed in the structure.

#### Minor helices

The most frequent minor helices are K′ (100%) and B′ (83%), followed by A ′ (63%), B″ (61%), K″ (46%), J′ (35%), L′ (30%), F′ (28%) and G′ (25%). Helices A′, B″, K″, L′ are very short, so it can often happen that they are not formed at all. On the other hand, the low occurrence of helices F′, G′, J′ can be explained by their absence in bacterial CYPs (roughly 2/3 of all structures) – for more details see section “Comparison of bacterial and eukaryotic CYPs “. Furthermore, the flexible F′G′-loop is often not modeled in the experimental structures.

#### Strands

In sheet β1, strands β1-1, β1-2, β1-3, β1-4 are always present, β1-5 is found in 96% of the structures, β1-0 in 44%. Strand β1-0 is less frequent because it is very short, and it is in the region of variable secondary structure before helix A. Sheet β2 is always present. In sheet β3, strands β3-2, β3-3 are present in more than 98% of structures (if they are missing, it is because the corresponding residues are not modeled in the experimental structure). β3-1 is sometimes not formed (present in 96%). Sheet β4 is found in 91% of structures – sometimes it is not formed. The remaining two sheets are much less frequent – β5 (25%) and β6 (46%) – which can be related to their very short length.

### Length of the SSEs

The typical length of individual helix classes varies substantially, from the minimal possible value of 3 residues (helices B″, L′) to 33 residues (helix I). On the other hand, β-strands in CYP structures are much shorter and range from 1 residue (strands β5-1, β5-2, β6-1, β6-2) to 5 residues on average (strands β1-1 through β1-4). The distribution of length of each SSE class is visualized by a violin plot in Fig. 3.

**Figure 3.**
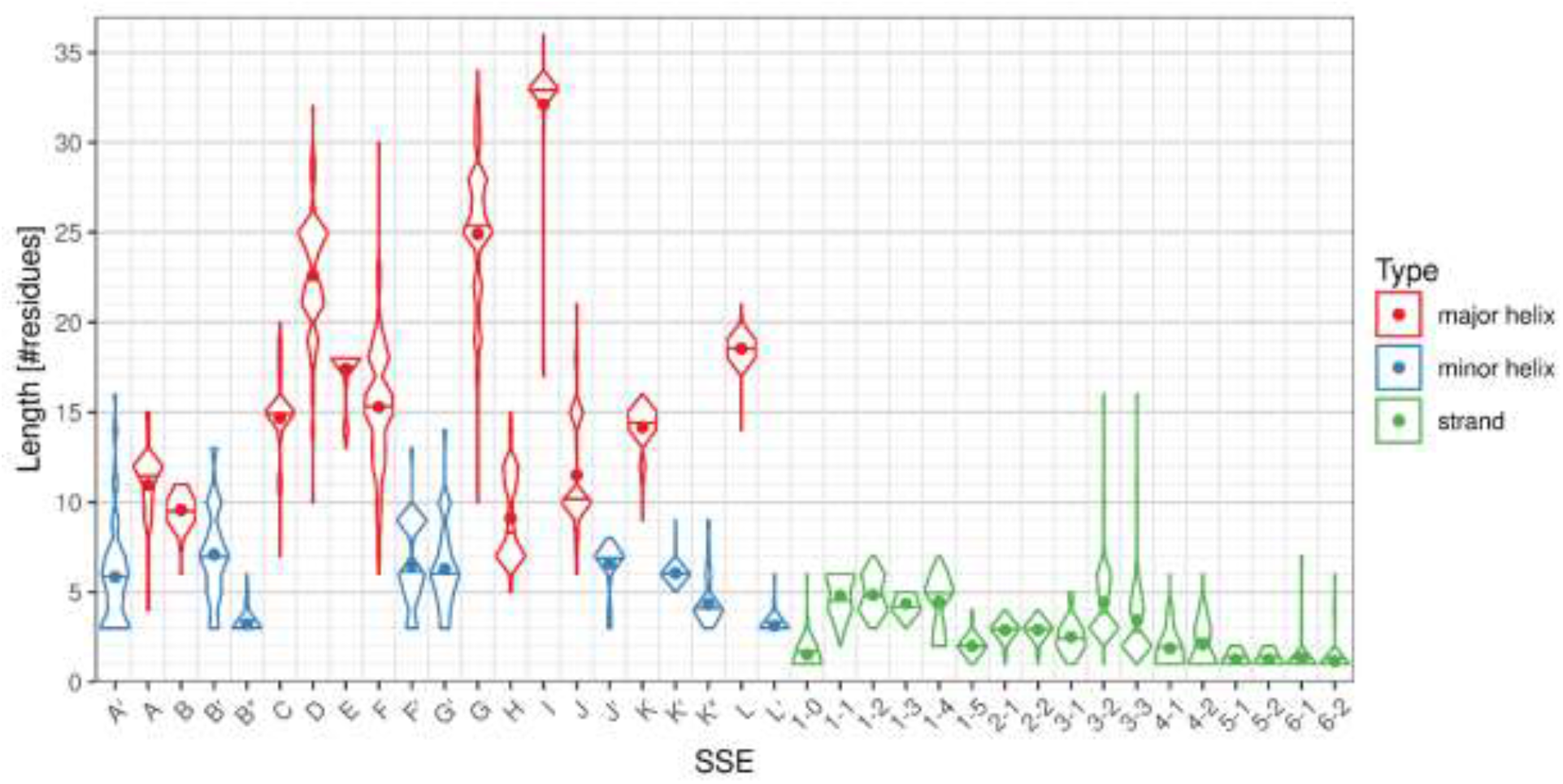
Distributions of length of individual SSE classes. The dots in the violin plot represent the mean, horizontal lines represent the median of the distribution. Non-existing SSEs are not included in the distribution.

#### Helices

The helix SSE classes differ not only by their average length but there are also great differences in the length variability. Helices with the most uniform length are the helices in the core of the structure (C, E, I, K, L), helices J′ and K′, and some very short helices on the edge of detection (B″, L′, if they exist, they are very short (3–4 residues), so there is no space for variability).

Helices with the most variable length are the helices in the regions of variable secondary structure (A′, A, B′, F, F′, G′, G, K″).

Some helices have a bimodal distribution:

- For helix J, there are two peaks in the length distribution at 10 and 15 residues, with almost no samples in-between. This is because the length is different in bacteria (10 residues) and eukaryotes (15 residues); within these groups the length is very uniform (for details see section “Comparison of bacterial and eukaryotic CYPs “).
- For helix H, there are two peaks at 7–8 residues and 11–12 residues (not as nicely separated as in the case of helix J). In this case it cannot be easily explained as in the case of helix J (although there is a preference for shorter length in bacteria). The difference between the peaks roughly corresponds to one turn of α-helix; the lengths between the two peaks must be unstable, because in such case the protein chain would have to continue in the opposite direction.
- Some other helices (e.g. D, F) also have complex length distri butions.

#### Strands

Sheet β1 has around 5 residues in each strand, with the exception of the last strand (β1-5), which has only 2 residues. Strand β1-0, if present, has 1–2 residues.

Sheet β2 has quite uniform length, in most cases 3 residues in each strand.

Sheet β3 has more variable length, because of the differences between bacterial and eukaryotic structures (for details see section “Comparison of bacterial and eukaryotic CYPs “). In both cases, the middle strand β3-2 is slightly longer, because it bridges β3-1 and β3-3.

Sheet β4 is quite short (usually 1–3 residues per strand).

The remaining sheets are very short – β5 usually contains 1–2 residues per strand, β6 only 1 residue per strand.

### Multiple sequence alignment and generic residue numbering in SSEs

For SSE classes with sufficient sequence conservation we created sequence logos (see Fig. 4) and selected the most conserved residue as the reference residue. Based on the reference residue, we established a generic residue numbering scheme similar to the schemes used for GPCRs (described by Isberg *et al*.^8^ and used throughout GPCRdb^36^) or immunoglobulins^19^.

**Figure 4.**
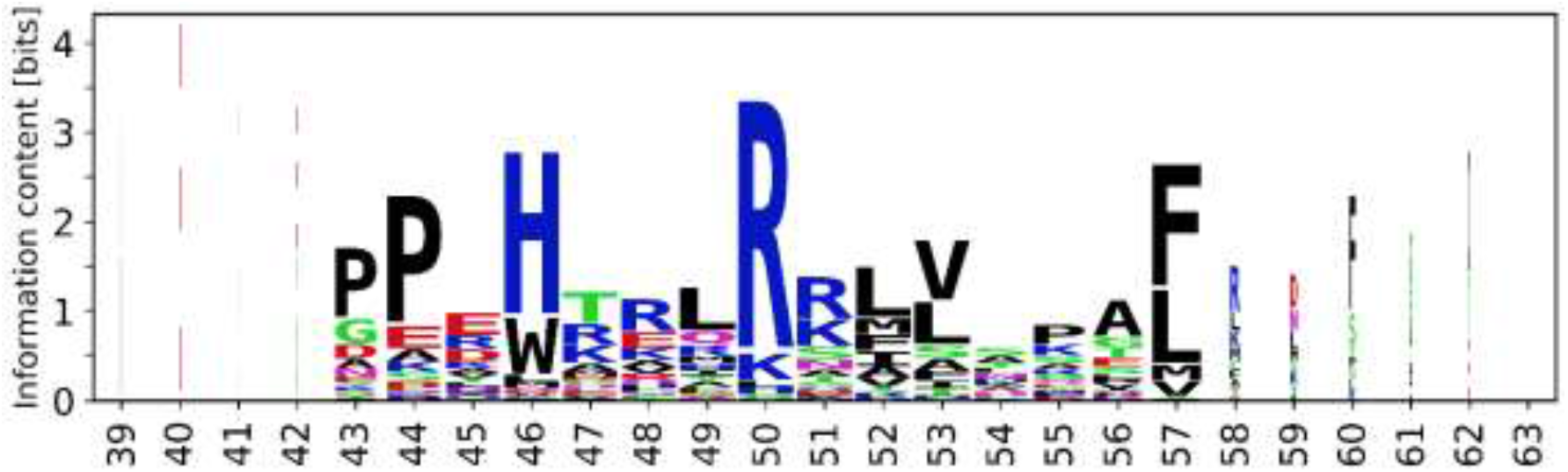
Sequence logo of helix C with generic numbering of residues. The remaining sequence logos are available in Supplementary Figures S3 and S4.

The reference residue is always numbered as @X.50, where X is the SSE label (character @ is added to avoid confusion in line notation). The remaining residues are then numbered correspondingly. An example of such residue identification is W120^@C.46^ in structure 2nnj (or generically @C.46), denoting the tryptophan residue 120 in helix C four positions before the reference residue, which is R124^@C.50^. Residue mutations can be also specified, e.g. W120A^@C.46^ (Fig. 4).Several examples of the usage of generic residue numbers can be found in section “Comparison of bacterial and eukaryotic CYPs”, demonstrating their usefulness.

We established the generic numbering for those SSEs, that contain at least one column in their logo with area (*c*_*i*_) greater than 2 bits: major helices B, C, E, H, I, J, K, L, minor helices J′, K′, K″, and strands β1-1, β1-2, β1-3, β1-4, β1-5, β2-2, β3-1, β3-2. Other SSE classes have insufficient sequence conservation and/or sequence alignment does not correspond to structure alignment (i.e. reference residues do not align in 3D). Therefore, it is impossible to establish a meaningful generic numbering for these SSE classes. All sequence logos are available in Supplementary Figures S3 and S4.

These logos are computed only from the dataset of the available structures (Set-NR) and thus will differ from logos obtained from the alignment of all available sequences. Still, when compared to Pfam representative proteome alignment (Supplementary Figure S5), they hold the key conserved residues and thus can be used for the definition of generic residue numbering.

### Comparison with substrate recognition sites

We compared the sequence variability of the SSEs with the positions of the substrate recognition sites (SRS) in CYPs reported by Gotoh^17^ and Zawaira *et al*.^18^ (see Table 1). From this comparison, we observe that most SRS sites (SRS1, SRS2, SRS3, SRS6) are located in the SSEs with the most variable sequence. CYPs are known for high substrate variability and Table 1 shows that the substrate recognition sites are mirrored in the structural variability of their SSEs.

**Table 1.** Comparison between the sequence conservation of the SSE classes (quantified by average information content) and the position of the substrate recogniti on sites from ref erences^17,18^. The SSEs are sorted from the most conserved to the most variable. The “Heme” column marks the SSEs with any atom within 8 Å from the heme cofactor.

| SSE label | Residue range (in 2nnj) | Heme | Average information content | Gotoh 1992 | Zawaira *et al.* 2011 |
| --- | --- | --- | --- | --- | --- |
| K'' | 409–412 |  | 2.18 |  |  |
| β1-5 | 96–97 | ✓ | 2.06 |  |  |
| β6-1 | 362 |  | 1.98 |  |  |
| K | 346–359 | ✓ | 1.98 | SRS 5 | SRS 5 |
| β6-2 | 477 |  | 1.87 |  |  |
| β3-1 | 164 |  | 1.84 |  |  |
| L' | 464–466 |  | 1.77 |  |  |
| L | 438–455 | ✓ | 1.76 |  |  |
| J | 317–331 |  | 1.76 |  |  |
| K' | 391–396 |  | 1.70 |  |  |
| J' | 339–345 |  | 1.67 |  |  |
| C | 117–131 |  | 1.67 |  | SRS 1 |
| β5-1 | 274 |  | 1.61 |  |  |
| β1-4 | 368–369 | ✓ | 1.60 | SRS 5 | SRS 5 |
| β3-3 | 456–459 |  | 1.55 |  |  |
| β2-1 | 374–376 |  | 1.55 |  |  |
| I | 284–316 | ✓ | 1.54 | SRS 4 | SRS 4 |
| β5-2 | 280 |  | 1.53 |  |  |
| β1-2 | 72–77 |  | 1.51 |  | SRS 1'b |
| β2-2 | 379–381 |  | 1.49 |  |  |
| G' | 220–226 |  | 1.47 |  | SRS 2 |
| B | 80–90 |  | 1.43 |  |  |
| β3-2 | 485–489 |  | 1.41 |  |  |
| F' | 211–219 |  | 1.38 |  | SRS 2 |
| H | 263–274 |  | 1.37 |  |  |
| β1-3 | 386–389 |  | 1.35 |  |  |
| β1-1 | 64–69 |  | 1.31 |  | SRS 1'b |
| E | 166–183 |  | 1.22 |  |  |
| β1-0 | 32 |  | 1.20 |  |  |
| D | 141–159 |  | 1.20 |  |  |
| B'' | 112–114 | ✓ | 1.20 |  |  |
| β4-2 | 478–479 |  | 1.18 |  | SRS 6 |
| A' | 42–44 |  | 1.18 |  | SRS 1'a |
| A | 50–61 |  | 1.12 |  |  |
| G | 227–254 |  | 1.04 | SRS 3 | SRS 3 |
| F | 192–209 |  | 0.86 | SRS 2 | SRS 2 |
| B' | 101–107 | ✓ | 0.73 | SRS 1 | SRS 1 |
| β4-1 | 473–474 |  | 0.70 | SRS 6 | SRS 6 |

Two exceptions to this observation are SRS4 and SRS5, located in the highly or moderately conserved SSEs. This can be explained by their proximity to the heme cofactor – stabilization of the heme requires highly conserved amino acids in the neighboring SSEs.

### Comparison of bacterial and eukaryotic CYPs

We compared the frequencies of occurrence of individual SSE classes between bacterial and eukaryotic structures (Fig. 5) as well as their distributions of length (Fig. 6).

**Figure 5.**
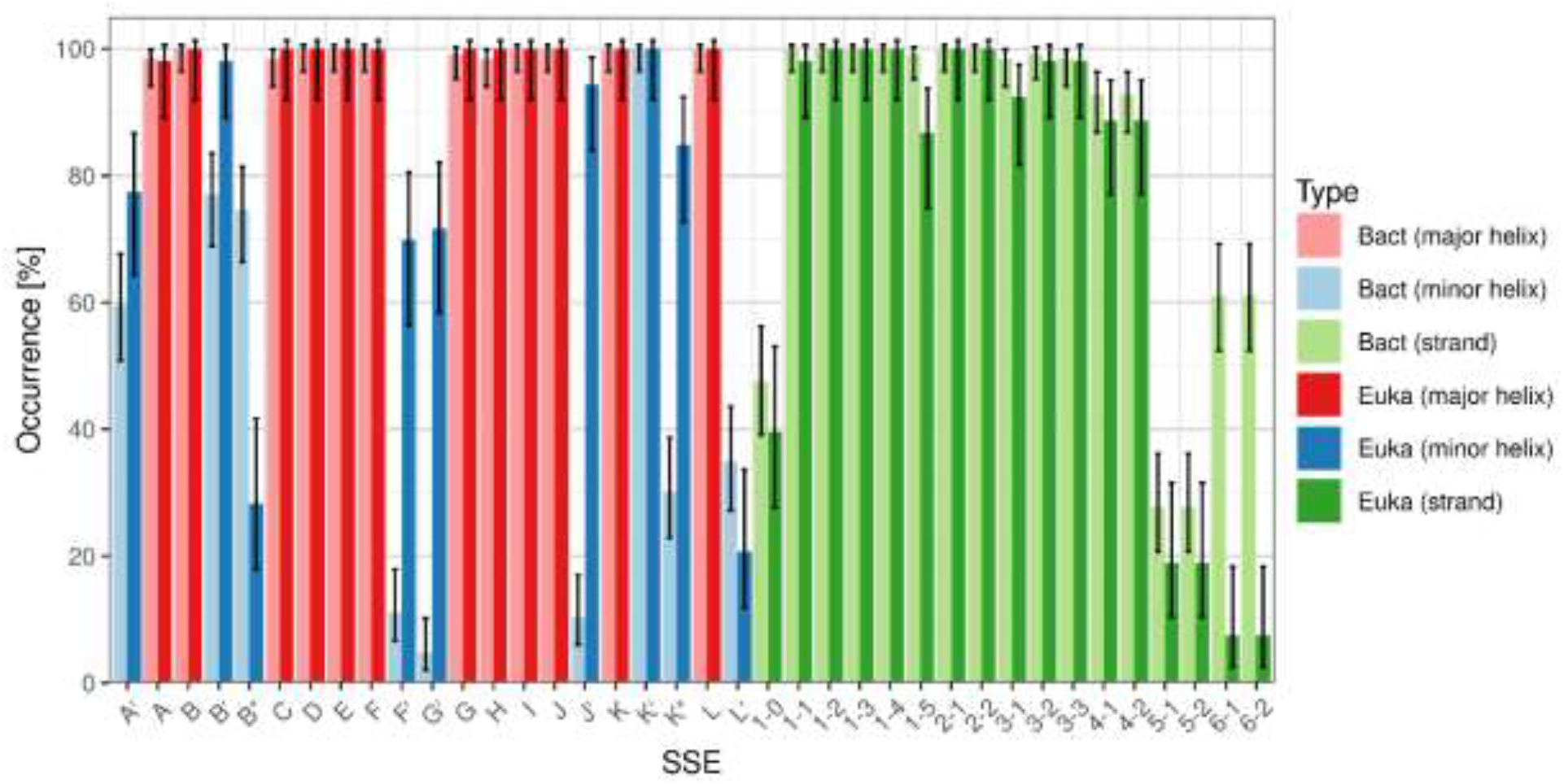
Comparison of SSE class occurrence in bacterial (Bact) and eukaryotic (Euka) CYP structures.

**Figure 6.**
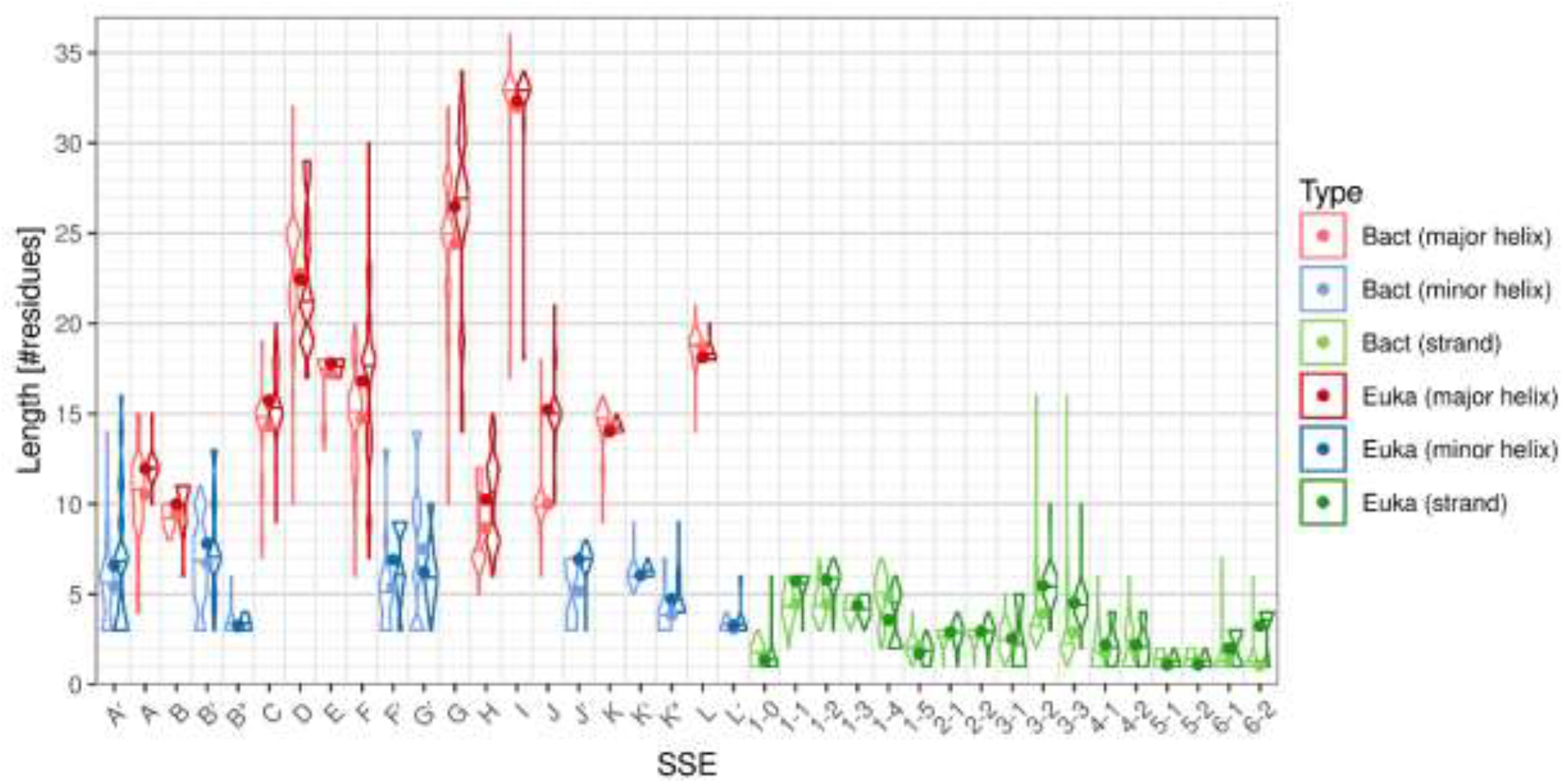
Comparison of SSE class length distribution in bacterial (Bact) and eukaryotic (Euka) CYP structures. The absent SSEs are not included in the distributions.

Helices A′, B′, F′, G′, J′ and K″ are significantly more frequent in eukaryotes. This is in agreement with literature^14,23^ reporting that F′ and G′ are typically not present in bacteria. However, sometimes a short helix can be formed between F and G in bacterial structures, which is then automatically annotated as F ′ or G′, so the observed occurrence is not equal to zero even in bacteria.

Conversely, helix B″, sheet β6 and strand β1-5 occur more frequently in bacteria.

Helices A, A ′, C, F, G, H, J, J′ and strands β1-1, β1-2, β3-2, β3-3, β6-2 are longer in eukaryotes, while strand β1-4 tends to be longer in bacteria. The case of β6-2 can be explained by the merging of β6-2 and β4-2 in some eukaryotic structures. The Kolmogorov-Smirnov test reports significant length difference for a few more SSE classes (B, K″ towards eukaryotes; D, K, L, β1-0, β1-5 towards bacteria), however in these cases the mean length differs by less than one residue.

Complete results of the statistical tests, including the *p*-values, can be found in Supplementary Tables S2 and S3.

A notable difference is visible in the region of J and J ′ helices. Their typical length in eukaryotes is 15 and 7 residues, respectively, while in bacteria helix J has typically 10 residues and J ′ is not present at all. However, there is an anomalous group of 5 bacterial CYPs whose helices J and J ′ have lengths 15 and 7 residues, exactly as observed in eukaryotic CYPs. This group includes:

- the heme domain of the flavocytochrome P450 BM3 (*Bacillus megaterium*, PDB IDs 3kx3, 6h1l and 6h1t (these map to different UniProt ID s but have the same sequence of the heme domain)), where helix J’ plays a role in the interaction with redox flavodomain,
- CYP 51 (*Mycobacterium tuberculosis*, PDB ID 2ci0),
- CYP 51 (*Methylococcus capsulatus*, PDB ID 6mcw),
- CYP 120A1 (*Synechocystis sp*., PDB ID 3ve3),
- CYP 170A1 (*Streptomyces coelicolor*, PDB ID 3dbg).

We further investigated this anomalous bacterial group and discovered more similarities to the eukaryotic structures:

- The sequence of their helix J resembles the sequence of eukaryotic helix J – most notably the glutamic acid E@J.59 (see section “Multiple sequence alignment and generic residue numbering in SSEs”) is highly conserved in both the eukaryotic CYPs (98%) and the anomalous group (100%) due to interactions with backbone amide groups of residues @K.40 and @K.41, while being much less conserved in the rest of the bacterial CYPs (7%), where this interaction is not observed.
- The region between helix K ′ and the heme binding site is approximately 9 residues longer in the eukaryotic and anomalous CYPs compared to the regular bacterial CYPs (roughly 39 residues in the eukaryotic and anomalous CYPs, 30 residues in the regular bacterial CYPs). Furthermore, in eukaryotic and anomalous CYPs, this region typically contains helix K ″ (in 85% of the eukaryotic and in all anomalous CYPs), which is usually not present in the regular bacterial CYPs (only in 26%).

These deviations are concentrated in the region which has been reported as the interface for binding of the redox partner (for P450 BM3^37^ and human mitochondrial CYP11A1^38^). This suggests that these deviations may be functionally related to the interaction with the redox partner.

Generally, the bacterial and mitochondrial eukaryotic CYPs receive electrons from a small iron-sulfur protein, while the microsomal eukaryotic CYPs receive electrons from a flavoprotein, like NADPH-cytochrome P450 oxidoreductase (CPR)^37^. Three of the five anomalous bacterial CYPs violate this interaction pattern : *Bacillus megaterium* P450 BM3 (aka CYP 102A1) contains flavodomain on C-terminus as a part of its sequence; *Mycobacterium tuberculosis* CYP 51 interacts with NADPH-hemoprotein reductase^39^; *Streptomyces coelicolor* CYP 170A1 is also known to be reduced by NADPH^40^. However, we have not found similar interactions with flavodomain for *Methylococcus capsulatus* CYP 51, but it belongs to a specific class containing Fe-4S ferredoxin-type on its C-terminus^41^. We have found no information about interactions with redox partners for putative *Synechocystis sp*. CYP 120A1, but from the anomalous motif we can hypothesize interactions with some NADPH-hemoprotein reductase.

In some other aspects the anomalous CYPs behave as typical bacterial CYPs – there is no F′ and G′ helix in the FG-loop; the A-propionate side chain of the heme is oriented to the distal side (towards the substrate binding pocket). We can therefore hypothesize that this group represents evolutionary transition towards eukaryotic CYPs – this is also supported by the fact that the anomalous bacterial CYPs group with the eukaryotic sequences in the phylogenetic tree from Set-NR (see Supplementary Fig. S6).

### SecStrAPI: how to get to our annotations

All annotations which are mentioned in this paper are publicly available through SecStrAPI^42^.

The annotations can be downloaded directly (in JSON format, described in detail on the website) or can be accessed through PyMOL plugin *secstrapi_plugin*.*py*, which is available on the website and serves for simple and quick visualization of the SSE annotations.

Any cytochrome P450 structure, including new structures not included in our dataset, can be uploaded and annotated in our web application SecStrAnnotator Online^43^.

## Limitations of the method

The presented methodology is in principle applicable to any protein family of interest. The workflow is almost fully automated – the bottleneck is the preparation of the annotation template. An appropriate template domain must be selected, and its annotation must be found in literature or created from scratch (especially in the families where no annotation conventions exist). However, we are currently developing software for automatic template generation, which will allow generalization of the annotation pipeline over all CATH protein families.

Another limitation is of course the fact that the inputs are experimental protein structures which are often incomplete. If an SSE is located in region which is not modeled in the experimental structure, then it will not be annotated and its observed occurrence in the family will be lower than its real occurrence. In the same way, this can affect the observed SSE length. This happens most often in the peripheral parts of the structure, in case of CYPs the FG-loop, BC-loop, HI-loop, JK-loop, sheet β3, and N-terminal part. The experimental setup (e.g. crystallization conditions, resolution, refinement procedure) can also induce small structural variations, which may influence the exact length of the detected SSEs.

The natural diversity of the structures within a family can make it difficult to find the correct SSE annotation. For example, when a region in the template protein is occupied by a single helix but the equivalent region in another protein contains two helices, it might not be clear which of the two helices should be annotated. SecStrAnnotator will base the annotation on optimization of the overall score, which takes into account purely the structural information. In extreme cases, this automatic annotation can be different from the annotation by an expert. We list the regions with the most uncertain annotation in section “Regions of variable secondary structure “. The annotation files contain the metric value for each SSE (higher values imply greater difference from the template and thus lower confidence of annotation).

A similar issue is related to the secondary structure assignment – an SSE can be so strongly deformed (kinked) that the two parts of the SSE will be assigned as two separate SSEs and only one part will be correctly annotated. This can be seen especially in the case of helix I (which is known to contain a kink^14^) – its typical length is 33 residues but in some structures we observe a length of 17–18 residues (i.e. only one part of the broken helix I).

Still all these complications are limited to a small number of marginal cases and they do not significantly affect the overall view of the family.

## Conclusions

We presented a workflow for description of the secondary structure anatomy of a protein structural family – automatic annotation of secondary structure elements, analysis of their frequency of occurrence, typical length, position, amino acid composition and the variability of these properties.

We demonstrated these methods in the case study of the Cytochromes P450 (CYP) family. The characteristic SSEs of the family are 14 helices A, B, B′, C, D, E, F, G, H, I, J, K, K ′, L and 4 sheets β1, β2, β3, β4, which occur nearly in all structures. Optional SSEs include helices A ′, B″, F′, G′, J′, K″, L′, sheets β5, β6, and strand β1-0. Some of these SSE classes are very uniform in length (the core helices C, E, I, K, L, but also J ′, K′), while some show extensive length variation (A′, A, B′, D, F, F′, G′, G, H, J, K″, β3-2, β3-3). The shortest helices B″, L′ and sheets β5, β6 are on the edge of detection.

For the SSE classes with sufficient sequence conservation we have established a generic residue numbering scheme (e.g. W120^@C.46^) similar to that used for the GPCR family. Unfortunately, this was not possible in the variable regions (A, B′, D, F, G), which are responsible for substrate uptake and substrate recognition.

We also compared the eukaryotic and bacterial members of the CYP family. The most substantial difference is the absence of helices F′, G′, J′ in bacteria. Helices A ′, B′, K″ are also rarer in bacteria, while B ″, β6 and β1-5 are more common in bacteria than in eukaryotes. Many SSEs tend to be longer in either eukaryotes (A, A ′, F, G, H, J, J′, β1-1, β1-2, β3-2, β3-3, β6-2) or bacteria (β1-4).

Strikingly, we also identified a small group of 5 bacterial CYPs with typical eukaryotic features in the region of helices J–L, which can be explained by the interaction with the redox partner.

Automatic annotation of CYP SSEs allows not only the orientation in the structure but also among its channels, which play a crucial role in substrate recognition and product egress.

All the utilized software tools and the obtained data, including secondary structure annotations and generic residue numbers, are available at our website^21^. The annotations can be easily visualized with a PyMOL plugin. CYP structures can be also annotated by SecStrAnnotator Online^43^. While the annotations are now only specific for the CYP family, the software is applicable in principle to all protein families with defined annotations. We are currently working on the generalization of the annotation pipeline over all CATH protein families.

## Methods

### Datasets

#### Set-NR

A list of protein domains annotated as Cytochrome P450 was obtained from SIFTS resource^44^ via PDBe REST API^45^ (/mappings endpoint) on 7 July 2020. More specifically, annotations originating from databases CATH and Pfam (accessions 1.10.630.10 and PF00067) were merged to obtain 1855 protein domains located in 1012 PDB entries. The information about residue ranges was discarded and whole chains were taken instead (this was necessary because Pfam often wrongly annotates only a small portion of the chain). The domains were mapped to UniProt IDs and the best-quality domain was selected for each UniProt ID. The quality was measured by “overall_quality” obtained from PDBe REST API^45^ (/validation/summary_quality_scores/entry endpoint). The domains which map to no UniProt ID were excluded. The resulting Set-NR (non-redundant) contains 183 protein domains.

#### Set-NR-Bact and Set-NR-Euka

Domains from Set-NR were mapped to their source organism using PDBe REST API^45^ (/pdb/entry/molecules endpoint) and divided into four subsets based on their superkingdom using NCBI Taxonomy^46^: Set-NR-Bact (Bacteria, 126 structures), Set-NR-Euka (Eukaryota, 53 structures), Set-NR-Arch (Archaea, 3 structures) and Set-NR-Viru (Viruses, 1 structure). However, Set-NR-Arch and Set-NR-Viru were not analyzed separately because of their small size.

All data, including the lists of PDB IDs and UniProt IDs for each dataset, are available in the Zenodo repository^47^.

### Template annotation

Since SecStrAnnotator requires an annotated template structure, we have chosen a template domain based on multiple selection criteria.

First, the template should contain all SSE classes. Thus, we considered only the eukaryotic structures (Set-NR-Euka).

Second, the template structure should be an “average” structure which is as similar to all the others as possible. Therefore, we compared each pair of structures in Set-NR-Euka by *cealign* command in PyMOL and calculated pairwise Q-scores^48^. Then for each structure we calculated the average Q-score against all the other structures *Q*_avg_. We selected the structure with the highest *Q*_avg_ as the template domain, which was 2nnjA (human CYP 2C8).

Third, template structure should have sufficient resolution, quality and should not contain unmodeled loops etc. The selected domain 2nnjA meets these criteria (resolution 2.28 Å, overall quality 42.46, observed residue range 10 –472 covers the whole region of interest).

Secondary structure annotation was mapped from the annotation of CY P 2C9 by Rowland *et al*.^10^ with several added SSE classes, as described in section “Traditional SSE nomenclature in the CYP family “ and shown in Fig. 1a. General methods for the selection and annotation of the template with and without any prior knowledge can be found our earlier publication^20^.

### Annotation procedure

The annotation was performed using our software SecStrAnnotator. The current version 2.2 has been improved since the original publication^20^ of SecStrAnnotator 1.0, the most significant changes being the support for mmCIF files, label_* numbering, revised secondary structure assignment method (geom-hbond2), and detection of structural irregularities within SSEs. Switching from .NET Framework to .NET Core enabled more consistent usage across operating systems. Furthermore, many additional scripts have been added, thus creating the SecStrAnnotator Suite and facilitating the automation of the whole analysis pipeline (including automatic selection of the non-redundant set, sequence alignment, generic residue numbering, sequence logo visualization, statistical evaluation, visualization through a PyMOL plugin). To overcome the need of installation, we also introduced the SecStrAnnotator Online^43^ and SecStrAPI^42^ with precomputed annotation results.

The annotation algorithm consists of three main steps: secondary structure assignment, structural alignment and SSE matching. Detailed description is provided in Midlik *et al*.^20^. SecStrAnnotator was run with these settings: --ssa geom-hbond2 --align cealign --matching mom --soft --maxmetric 25,0.5,0.5 --label2auth --verbose.

### Statistical evaluation

All statistical tests were performed in R (version 3.4.4-1ubuntu1) using the *stats* library. The plots were generated in R using the *ggplot2* library.

#### Comparison of bacterial and eukaryotic dataset

The occurrence of each SSE class in Set-NR-Bact and Set-NR-Euka was modeled as a binomial distribution, and the two datasets were compared by the test of equal proportions (*prop*.*test*) with *α* = 0.05.

The distribution of length (number of residues) of each SSE class was compared between Set-NR-Bact and Set-NR-Euka. Where the medians of the eukaryotic and bacterial distribution were not equal, the two-sample Kolmogorov-Smirnov test (*ks*.*test*) with *α* = 0.05 was used to decide if the difference between the distributions is significant. Non-existing SSEs were not included in the length distributions.

### Multiple sequence alignment

The amino acid sequences of the individual SSE classes were extracted from Set-NR and aligned using an in-house algorithm NoGapAligner which allows gaps only at the beginning and at the end but not within a sequence (this is necessary in order to establish generic residue numbering). Substitution matrix was BLOSUM62 and the gap penalty was set to 10. Sequence logos were rendered using *logomaker* module for Python^49^.

For every position *i* in each multiple sequence alignment, the information content *R*_*i*_ and the conservation measure *c*_*i*_ were calculated as follows:

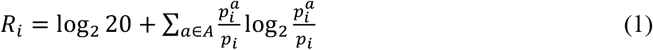

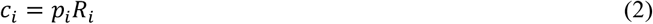

where *A* is the set of 20 standard amino acids, *pi*^*a*^ is the fraction of sequences having amino acid *a* at position *i, p*_*i*_ is the fraction of sequences having any amino acid (not a gap) at position *i*^50^. In the visual form of a logo, *p*_*i*_ and *R*_*i*_ correspond to the width and height of the *i*-th column, thus *c*_*i*_ corresponds to the area of the column. *R*_*i*_ is expressed in bits and its values can range from 0, for a position with 20 equiprobable amino acids, to approximately 4.3 (log _2_ 20), for a position with one perfectly conserved amino acid. The position with the greatest *c*_*i*_ within the alignment was selected as the reference residue of the SSE class.

To be able to compare the overall conservation of individual SSE classes, we computed the average information content *R*_avg_ (i.e. average column height) of each logo as:

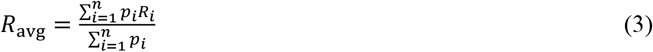

where *n* is the number of positions in the logo.

## Data availability

The datasets generated and analyzed during the current study are available in the Zenodo repository, https://doi.org/10.5281/zenodo.3939133^47^.

## Acknowledgements

This work was supported by the Ministry of Education, Youth and Sports of the Czech Republic under the project CEITEC 2020 [LQ1601]; ELIXIR-CZ research infrastructure project including access to computing and storage facilities [LM2018131]; European Regional Development Fund – projects ELIXIR-CZ [CZ.02.1.01/0.0/0.0/16_013/0001777]. A.M. was also supported by Brno Ph.D. Talent Scholarship funded by Brno City Municipality. We would like to thank Veronika Bendová for her valuable advice on the statistical procedures.

## Author contributions statement

R.S., K.B., and J.K. conceived, designed, and cooperated the study. A .M. created the software and analyzed data. T.R.M. tested the software and performed sequence analyses. A .M., V.N., K.B., and R.S. interpreted the results. A.M., R.S., and K.B. wrote the manuscript. All authors reviewed and approved the final manuscript.

## Additional information

### Supplementary information

Supplementary information contains these parts: i) Supplementary note about detected structural irregularities associated with their most typical location (Supplementary Figure S1) as well as their occurrence (Supplementary Figure S2); ii) Sequence logos of individual SSEs together with generic residue numbering (Supplementary Figures S3-S5); iii) Phylogenic tree of CYP family sequences with structures (Supplementary Figure S6); iv) Supplementary tables with the definition of typical residue ranges on template (Supplementary Table S1), with comparison of SSE occurrences and lengths in bacterial and eukaryotic CYP structures (Supplementaty Table S2 and S3).

