## Supplementary Information for "Uncovering of cytochrome P450 anatomy by SecStrAnnotator"

### SUPPLEMENTARY INFORMATION
### Uncovering of cytochrome P450 anatomy by SecStrAnnotator

Adam Midlik[1,2], Veronika Navrátilová[3], Taraka Ramji Moturu[1,2], Jaroslav Koča[1,2], Radka Svobodová[1,2,*], Karel Berka[3,*]

[1] CEITEC – Central European Institute of Technology, Masaryk University, Brno 625 00, Czech Republic
[2] National Centre for Biomolecular Research, Faculty of Science, Masaryk University, Brno 625 00, Czech Republic
[3] Department of Physical Chemistry, Faculty of Science, Palacký University, Olomouc 771 46, Czech Republic

* Corresponding authors


### Supplementary Note: Structural irregularities

#### Introduction

Helices found in protein structures are traditionally distinguished into three types: $3_{10}$-helix, α-helix and π-helix, characterized by repetitive $i+3 \rightarrow i$, $i+4 \rightarrow i$ and $i+5 \rightarrow i$ backbone hydrogen bonds, respectively. However, the $3_{10}$ and π-helices are much less common than the α-helix and rarely span more than a few residues. Combinations of the hydrogen bonding patterns commonly occur in a single helical segment, such as $3_{10}$-α-$3_{10}$ or α-π-α[1]. Therefore, we can understand the α-helix as the standard pattern and the $3_{10}$ and π-helices as structural irregularities within this pattern.

A β-bulge is a region of irregularity in a β-sheet formed by two or more residues on one strand (long side) opposite a single residue on the other strand (short side). β-bulges are relatively frequent (on average two instances per protein) and occur primarily between antiparallel strands[2,3].

#### Detection of structural irregularities

The traditional (DSSP) distinction of helix types is based on the type of hydrogen bonds stabilizing the helix (type $3_{10}$: $i+3 \rightarrow i$, type α: $i+4 \rightarrow i$, type π: $i+5 \rightarrow i$ bonds). A DSSP helix is detected when there are at least two consecutive hydrogen bonds of the same type[4].

SecStrAnnotator uses a method for helix detection which focuses on the geometry of the protein backbone and allows abstraction from these hydrogen bonding patterns. However, it also reports the hydrogen bonds found in each helix (when run with --verbose).

The *contained types* of such helix are then determined by the occurrence of two consecutive hydrogen bonds of the same type (i.e. a DSSP helix) within the helix. A helix may contain $3_{10}$, α, π, or any combination of these types (helices not containing any type are rejected).

It should be kept in mind that all obtained results are based on the DSSP definition of a hydrogen bond, which is approximate and quite benevolent.

#### Occurrences of irregularities

##### *Beta-bulges*

We analysed the frequency of occurrence of β-bulges in individual β-sheets and found out that they are not distributed randomly but occur mostly in sheets β3 and β4 (see Supplementary Fig. S1). The most common are:

- classic β-bulge on sheet β4, with the long side in β4-2 and the short side in β4-1 (in 20.8% of the structures)
- classic β-bulge on sheet β3, with the long side in β3-3 and the short side in β3-2 (in 8.2% of the structures)

Bulges of other types occur rarely (less than 5% structures for each type). Bulges in sheet β1 occur in less than 5% structures; they are never found in sheets β2, β5, β6.

The β-bulges are much more common in the bacterial than the eukaryotic structures. Namely, the bulge on sheet β4 is found in 28.6% bacterial and 1.9% eukaryotic structures; the bulge on sheet β3 is found in 8.7% bacterial and 3.8% eukaryotic structures. In archaeal structures, sheets β3 and β4 are usually merged into a single sheet containing two or more bulges.

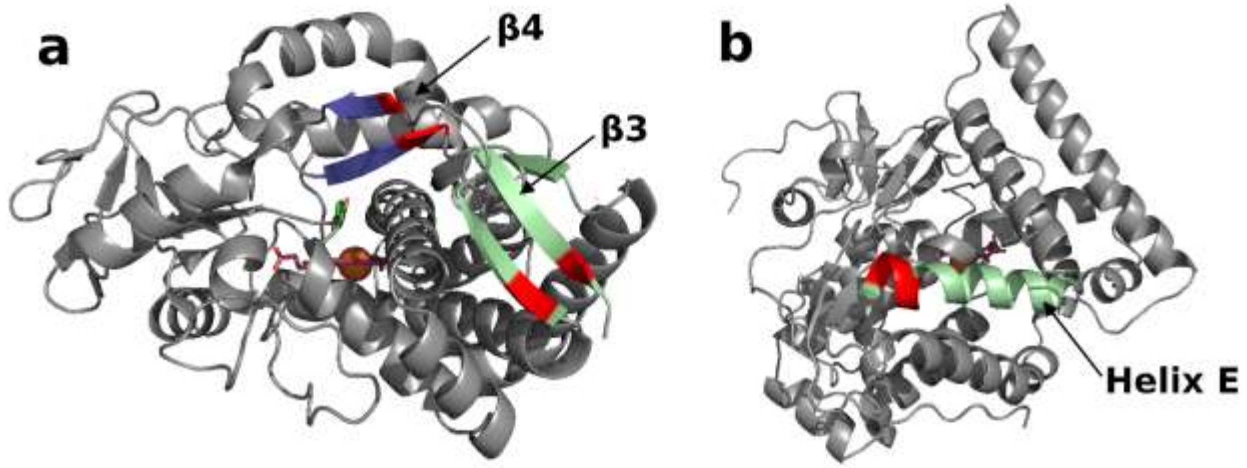


**Supplementary Figure S1. Location of the most common structural irregularities.** (**a**) The β-bulges in the bacterial CYP199A2 (PDB ID 4dnj); sheet β3 is shown in green, sheet β4 in blue, the bulges are highlighted in red. (**b**) The π-helix within helix E in the bacterial CYP142A2 (PDB ID 4uax); helix E is shown in green, the π-helix is highlighted in red.

##### $3_{10}$*-helices and* $\pi$*-helices*

A helix may consist of a single helix type ($3_{10}$, α, π) or may contain any combination of these basic types. We studied how often each of these types occurs in individual annotated helices.

The α-helix is of course the most abundant type and is present in 86.4% of all studied helices.

43.3% of all studied helices contain a $3_{10}$-helix. Most of these $3_{10}$-helices occur in the shortest minor helices, which are typically pure $3_{10}$ (L′, B″, K″), followed by J′, K′, and G′. Major helices with the highest content of $3_{10}$-helices are C, D, and F. In contrast, helices with the lowest occurrence of $3_{10}$-helical parts are L, E, and J (under 10%).

The π-helices are far less abundant than $3_{10}$-helices – only 6.3% of all helices contain a π-helix. The π-helices very often occur as a part of helix E (in 65.0% cases), followed by helix B (12.0%), helix I (8.2%), and helix B′ (5.3%). In other helices, their occurrence is under 5% (see Supplementary Fig. S2).

It is an interesting discovery that in 65.0% structures helix E contains a π-helix, and this fact might be related to the function or stability of the structures. This π-helix is typically located near the N-terminus of helix E (see Supplementary Fig. S1), and its occurrence is much higher in bacteria (87.3%) than in eukaryotes (9.4%). In the case of helix B this tendency is reversed (41.5% in eukaryotes, 0.0% in bacteria).

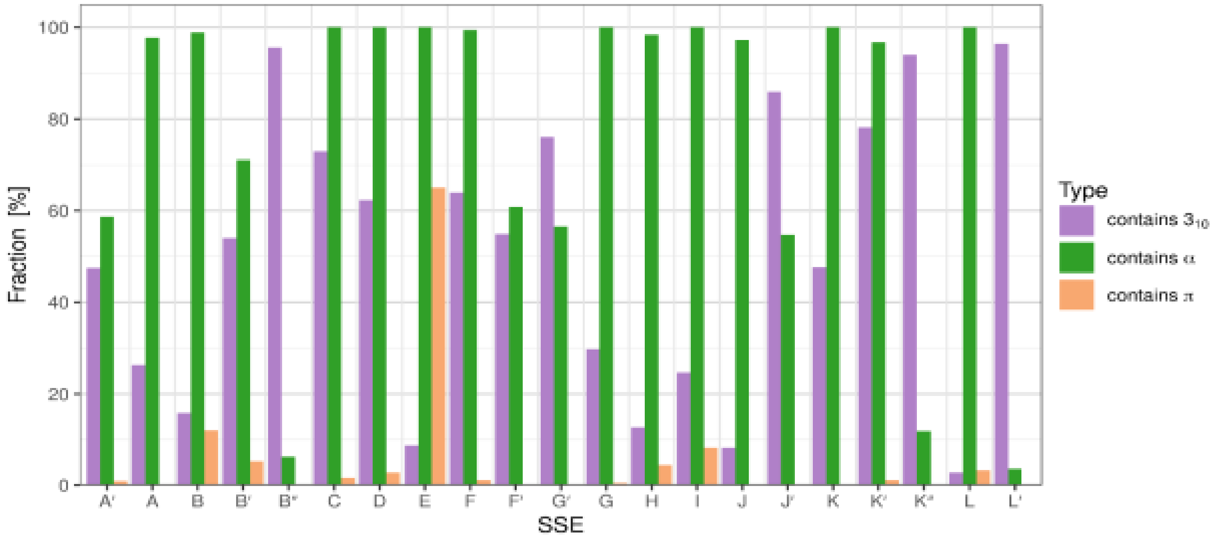


**Supplementary Figure S2. Percentage of helix types – $3_{10}$, α and π.**

### Sequence logos

#### Helices

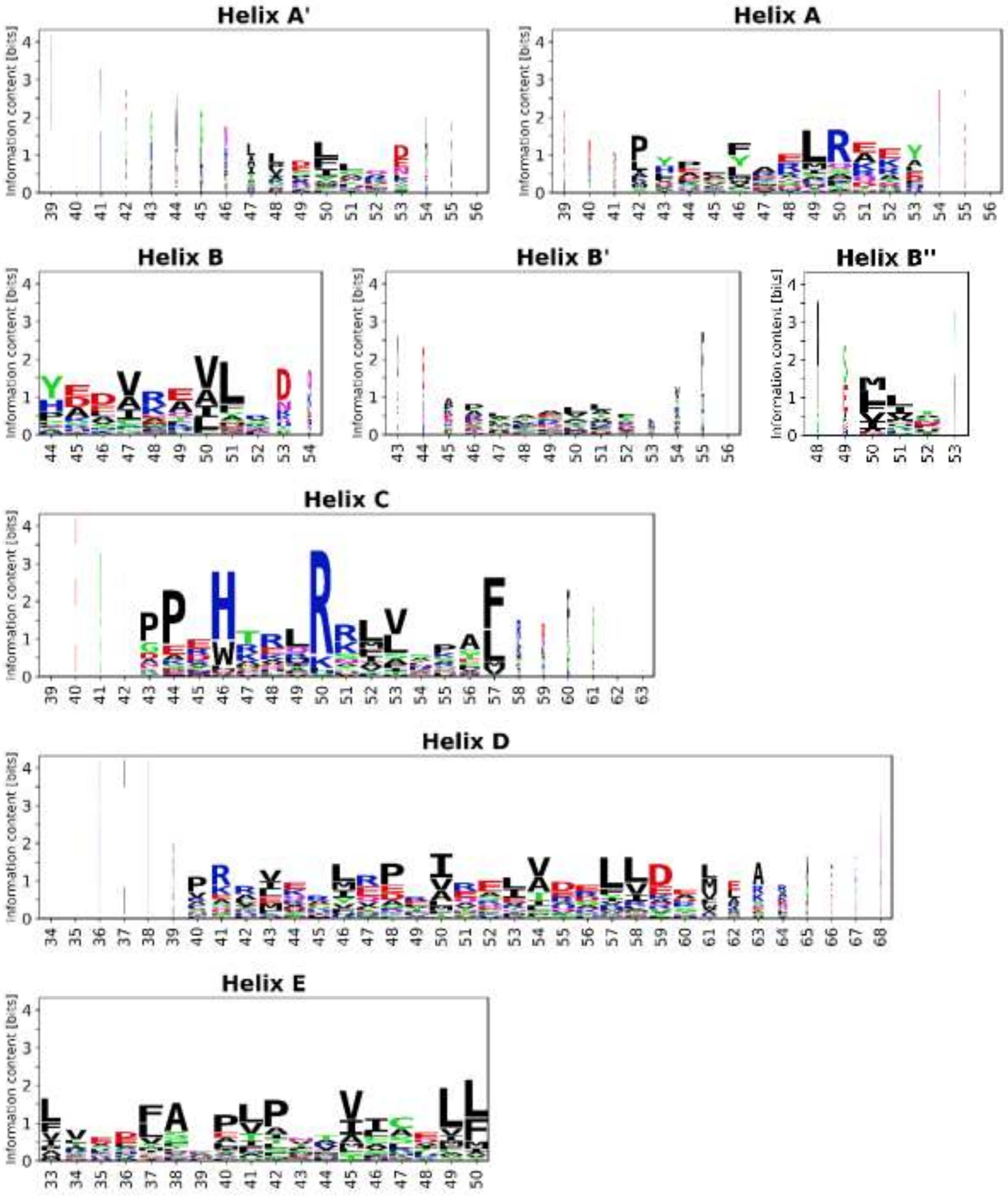


**Supplementary Figure S3. Sequence logos for the helices. (**Continues on the next page.)

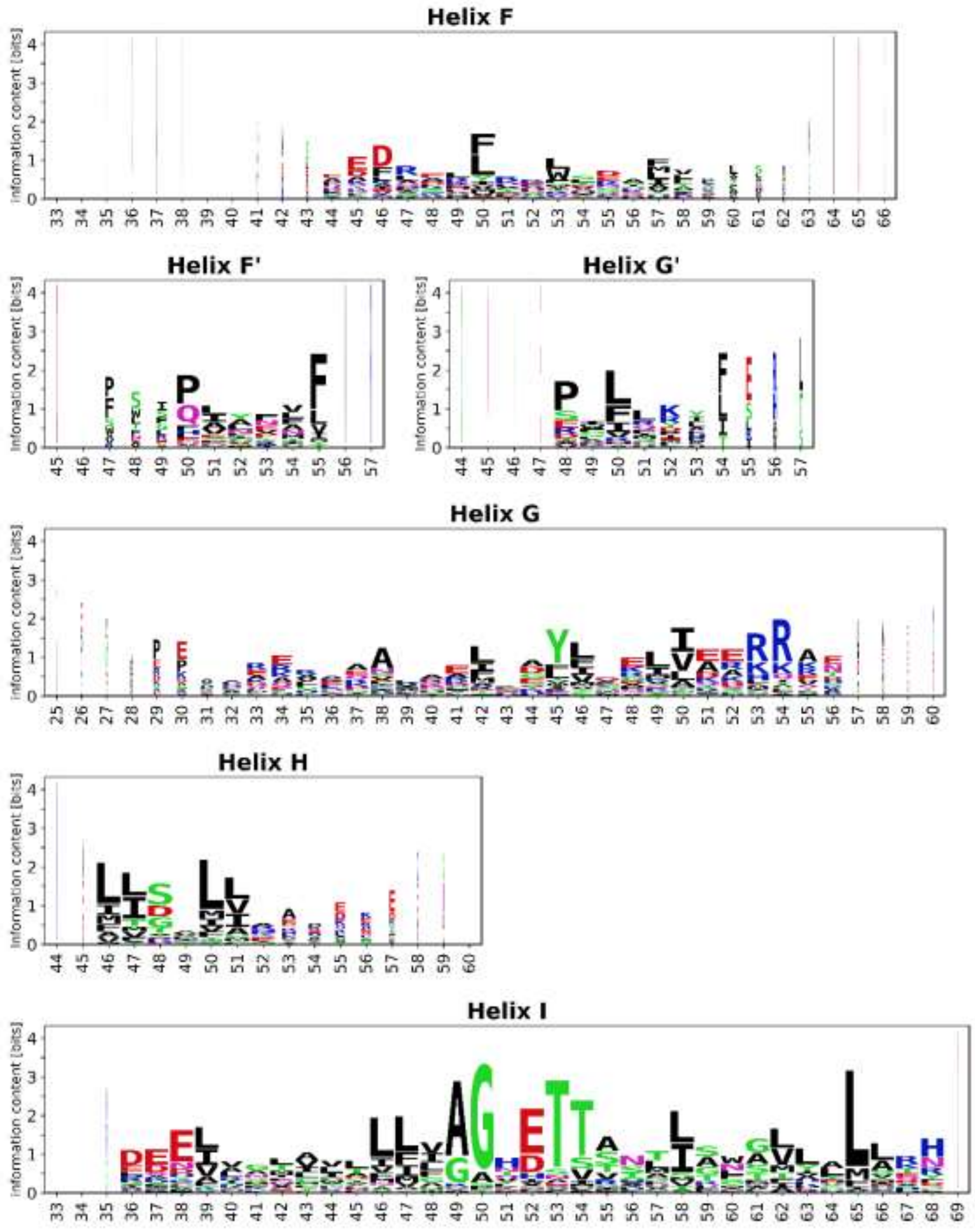


**Supplementary Figure S3 (continued). Sequence logos for the helices**. (Continues on the next page.)

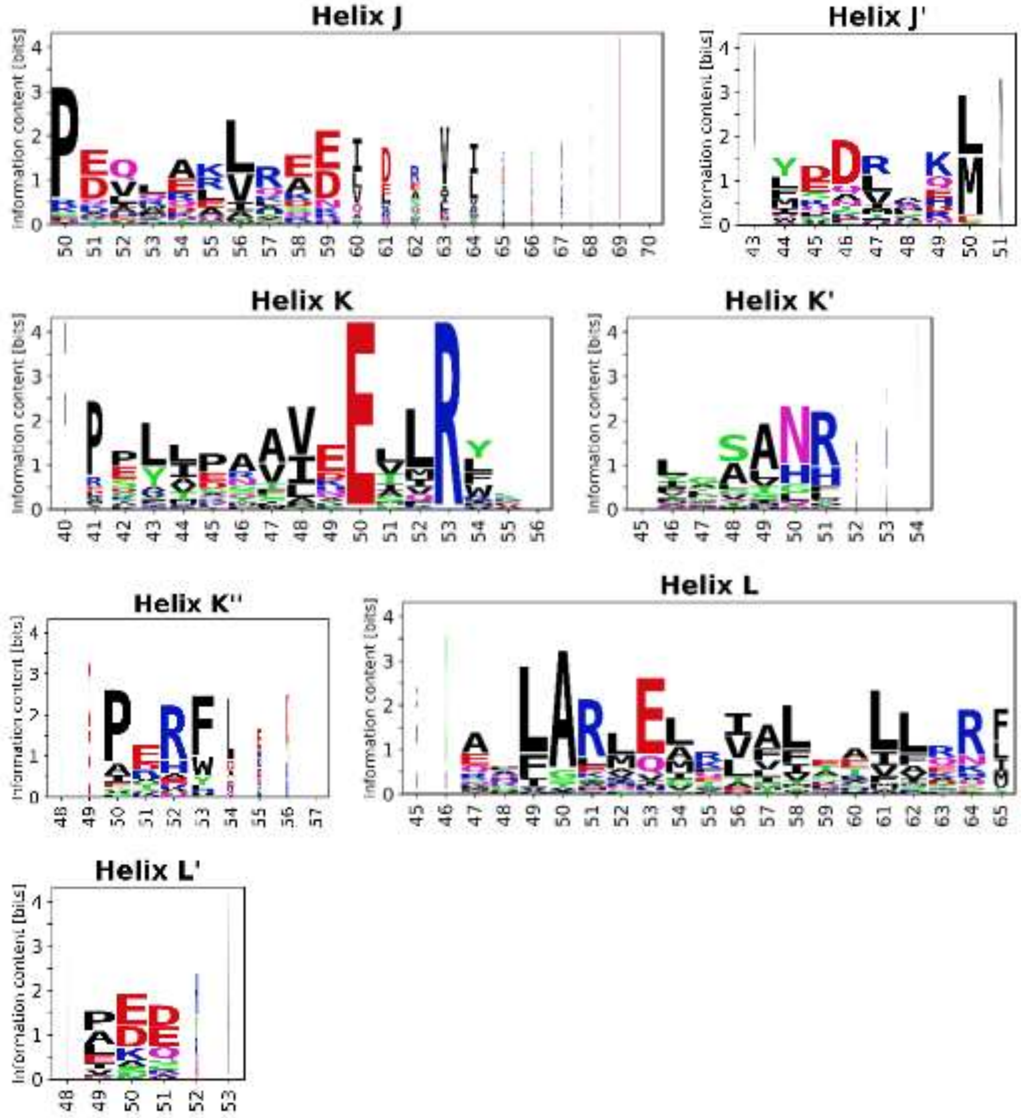


**Supplementary Figure S3 (continued). Sequence logos for the helices.**

#### Strands

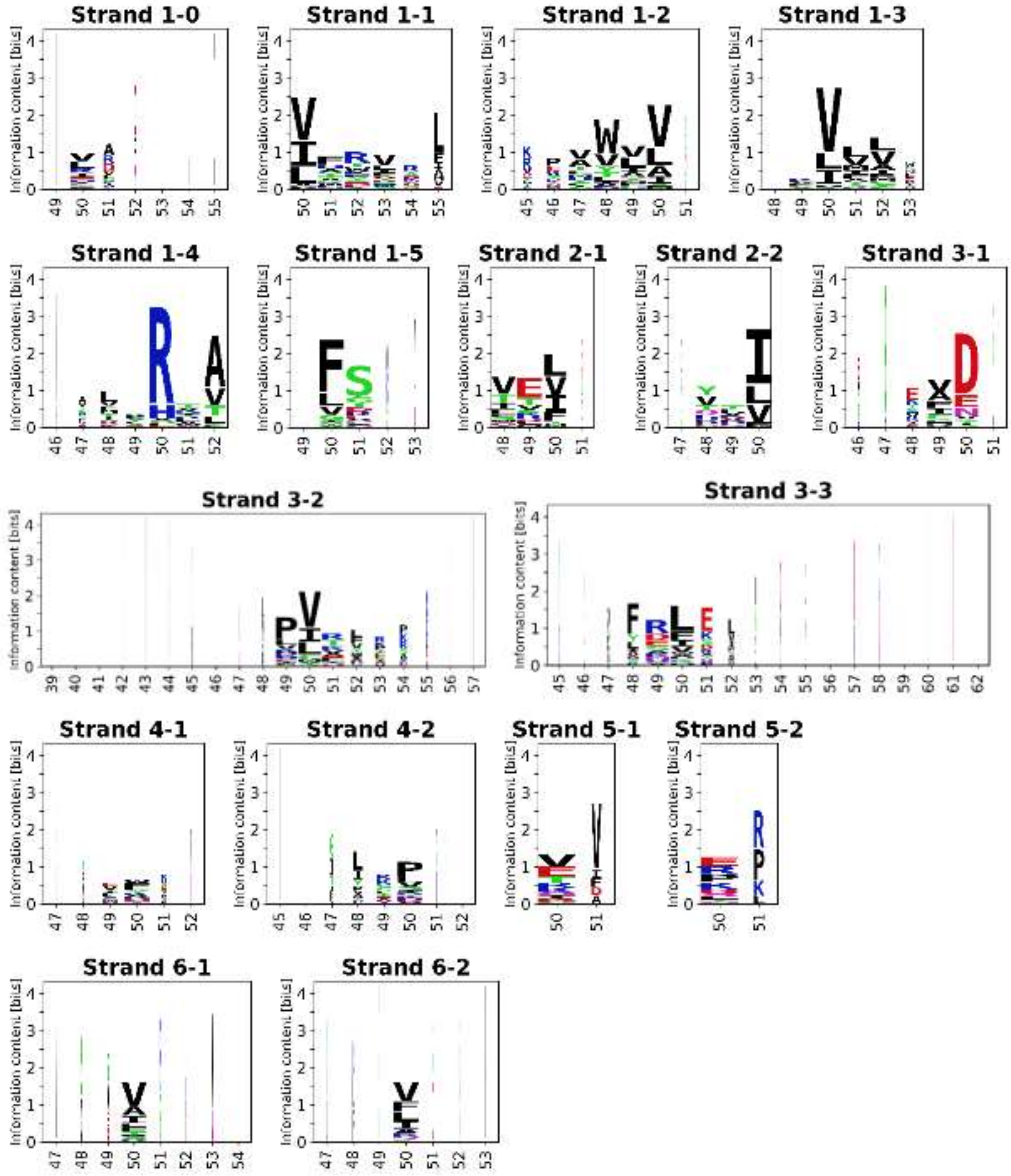


**Supplementary Figure S4. Sequence logos for the β-strands.**

#### Comparison of different approaches for sequence logo generation

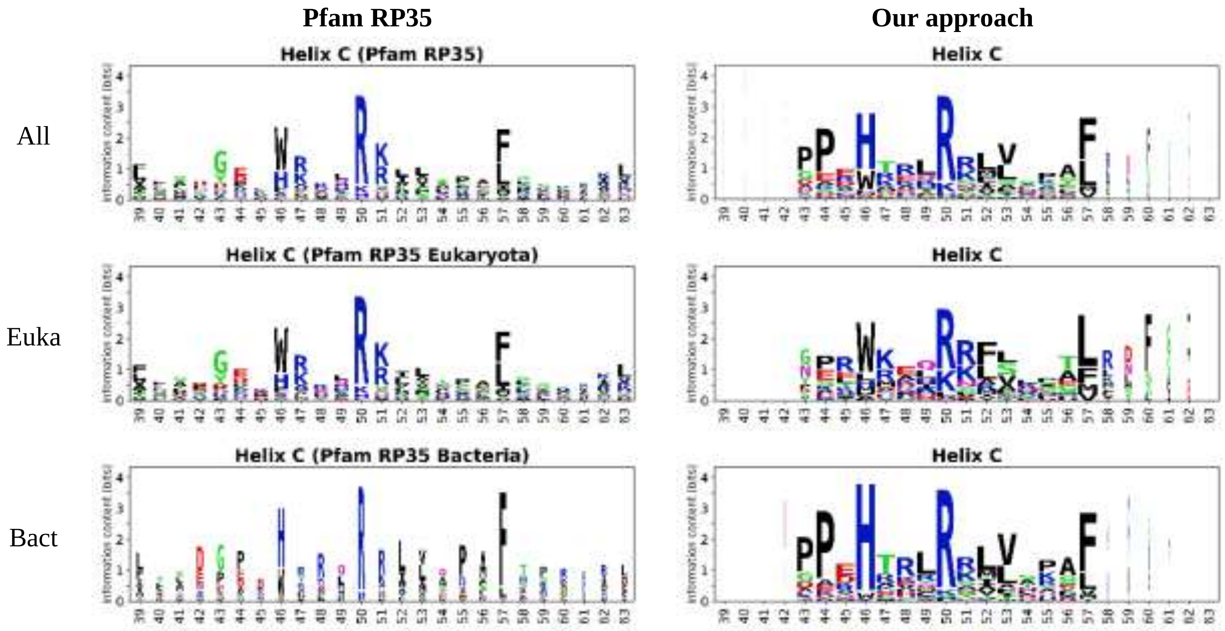


**Supplementary Figure S5. Comparison of sequence logo for helix C computed from Pfam representative proteome 35% sequence alignment and from Set-NR.**

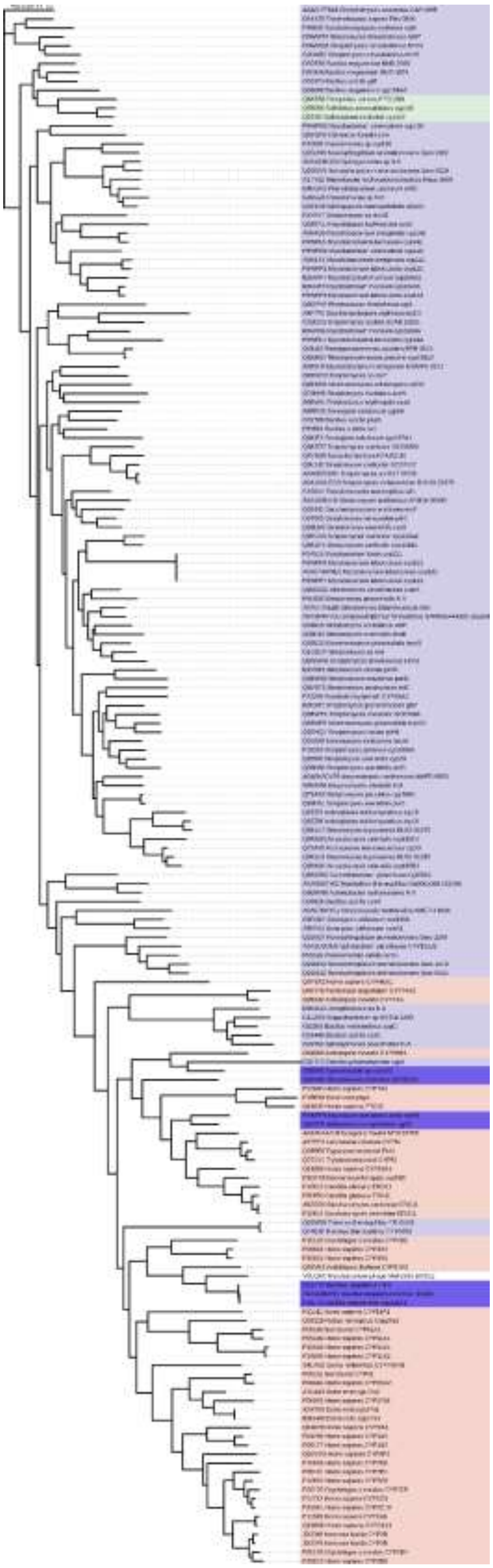

#### Phylogenic tree of CYP family structures

**Supplementary Figure S6. Phylogenetic tree based on multiple sequence alignment of full sequences from Set-NR.** Eukaryotic sequences are highlighted in red, bacterial in light blue, anomalous bacterial group in dark blue, archaeal in green, viral in white.

#### Supplementary Tables

**Supplementary Table S1.** Residue ranges of the SSEs in the PDB entry 2nnj (template annotation)

| SSE label | Residue range | SSE group |
|---|---|---|
| A′ | 42–44 | minor helix |
| A | 50–61 | major helix |
| B | 80–90 | major helix |
| B′ | 101–107 | minor helix |
| B″ | 112–114 | minor helix |
| C | 117–131 | major helix |
| D | 141–159 | major helix |
| E | 166–183 | major helix |
| F | 192–209 | major helix |
| F′ | 211–219 | minor helix |
| G′ | 220–226 | minor helix |
| G | 227–254 | major helix |
| H | 263–274 | major helix |
| I | 284–316 | major helix |
| J | 317–331 | major helix |
| J′ | 339–345 | minor helix |
| K | 346–359 | major helix |
| K′ | 391–396 | minor helix |
| K″ | 409–412 | minor helix |
| L | 438–455 | major helix |
| L′ | 464–466 | minor helix |
| β1-0 | 32 | strand |
| β1-1 | 64–69 | strand |
| β1-2 | 72–77 | strand |
| β1-3 | 386–389 | strand |
| β1-4 | 368–369 | strand |
| β1-5 | 96–97 | strand |
| β2-1 | 374–376 | strand |
| β2-2 | 379–381 | strand |
| β3-1 | 164 | strand |
| β3-2 | 485–489 | strand |
| β3-3 | 456–459 | strand |
| β4-1 | 473–474 | strand |
| β4-2 | 478–479 | strand |
| β5-1 | 274 | strand |
| β5-2 | 280 | strand |
| β6-1 | 362 | strand |
| β6-2 | 477 | strand |

**Supplementary Table S2. Comparison of the SSE occurrences in the bacterial and eukaryotic CYP structures.** Results of the test of equal proportions comparing SSE occurrences in Set-NR-Bact vs Set-NR-Euka. The column "Comparison" marks the significant differences at confidence level $\alpha$ = 0.05 (>: higher occurrence in Bacteria, <: higher occurrence in Eukaryota).

| SSE label | Occurrence (Set-NR-Bact) | Comparison | Occurrence (Set-NR-Euka) | Occurrence difference | *p*-value |
|---|---|---|---|---|---|
| A′ | 0.595 | < | 0.774 | 0.180 | 0.035 |
| A | 0.984 | | 0.981 | -0.003 | 1 |
| B | 1.000 | | 1.000 | 0.000 | NaN |
| B′ | 0.770 | < | 0.981 | 0.210 | 0.0012 |
| B″ | 0.746 | > | 0.283 | -0.460 | 1.8E-08 |
| C | 0.984 | | 1.000 | 0.016 | 0.89 |
| D | 1.000 | | 1.000 | 0.000 | NaN |
| E | 1.000 | | 1.000 | 0.000 | NaN |
| F | 1.000 | | 1.000 | 0.000 | NaN |
| F′ | 0.111 | < | 0.698 | 0.590 | 8.4E-15 |
| G′ | 0.048 | < | 0.717 | 0.670 | 1.3E-20 |
| G | 0.992 | | 1.000 | 0.008 | 1 |
| H | 0.984 | | 1.000 | 0.016 | 0.89 |
| I | 1.000 | | 1.000 | 0.000 | NaN |
| J | 1.000 | | 1.000 | 0.000 | NaN |
| J′ | 0.103 | < | 0.943 | 0.840 | 3.9E-26 |
| K | 1.000 | | 1.000 | 0.000 | NaN |
| K′ | 1.000 | | 1.000 | 0.000 | NaN |
| K″ | 0.302 | < | 0.849 | 0.550 | 6.1E-11 |
| L | 1.000 | | 1.000 | 0.000 | NaN |
| L′ | 0.349 | | 0.208 | -0.140 | 0.09 |
| β1-0 | 0.476 | | 0.396 | -0.080 | 0.41 |
| β1-1 | 1.000 | | 0.981 | -0.019 | 0.65 |
| β1-2 | 1.000 | | 1.000 | 0.000 | NaN |
| β1-3 | 1.000 | | 1.000 | 0.000 | NaN |
| β1-4 | 1.000 | | 1.000 | 0.000 | NaN |
| β1-5 | 0.992 | > | 0.868 | -0.120 | 0.0011 |
| β2-1 | 1.000 | | 1.000 | 0.000 | NaN |
| β2-2 | 1.000 | | 1.000 | 0.000 | NaN |
| β3-1 | 0.984 | | 0.925 | -0.060 | 0.12 |
| β3-2 | 0.992 | | 0.981 | -0.011 | 1 |
| β3-3 | 0.984 | | 0.981 | -0.003 | 1 |
| β4-1 | 0.929 | | 0.887 | -0.042 | 0.53 |
| β4-2 | 0.929 | | 0.887 | -0.042 | 0.53 |
| β5-1 | 0.278 | | 0.189 | -0.089 | 0.29 |
| β5-2 | 0.278 | | 0.189 | -0.089 | 0.29 |
| β6-1 | 0.611 | > | 0.075 | -0.540 | 1.5E-10 |
| β6-2 | 0.611 | > | 0.075 | -0.540 | 1.5E-10 |

**Supplementary Table S3. Comparison of the SSE length distributions in the bacterial and eukaryotic CYP structures.** Results of the Kolmogorov-Smirnov test comparing the SSE length distributions in Set-NR-Bact vs Set-NR-Euka. The column "Comparison" marks the significant differences at confidence level $\alpha = 0.05$ (>: longer in Bacteria, <: longer in Eukaryota). The column "*p*-value" contains the *p*-values for two-sided alternative hypothesis. Columns "$p_g$" and "$p_l$" contain values for one-sided alternative hypotheses.

| SSE label | Mean length (Set-NR-Bact) | Comparison | Mean length (Set-NR-Euka) | Mean difference | Median difference | *p*-value | $p_g$ | $p_l$ |
|---|---|---|---|---|---|---|---|---|
| A′ | 5.4 | < | 6.6 | 1.1 | 1.0 | 0.0049 | 0.77 | 0.0025 |
| A | 11.0 | < | 12.0 | 1.4 | 1.0 | 4.2E-11 | 0.41 | 2.1E-11 |
| B | 9.4 | < | 10.0 | 0.6 | 2.0 | 1.8E-09 | 0.79 | 9E-10 |
| B′ | 6.7 | | 7.8 | 1.1 | 0.0 | 0.05 | 0.96 | 0.025 |
| B″ | 3.2 | | 3.3 | 0.1 | 0.0 | 0.96 | 0.9 | 0.61 |
| C | 14.0 | < | 16.0 | 1.4 | 0.0 | 5.9E-05 | 1 | 3E-05 |
| D | 23.0 | > | 22.0 | -0.3 | -3.0 | 0.022 | 0.011 | 0.074 |
| E | 17.0 | | 18.0 | 0.6 | 0.0 | 0.17 | 1 | 0.083 |
| F | 15.0 | < | 17.0 | 2.1 | 3.0 | 1.7E-07 | 0.75 | 8.6E-08 |
| F′ | 5.4 | | 6.9 | 1.5 | 1.0 | 0.078 | 0.9 | 0.039 |
| G′ | 7.5 | | 6.2 | -1.3 | -1.5 | 0.39 | 0.2 | 0.38 |
| G | 24.0 | < | 26.0 | 2.1 | 2.0 | 1.3E-06 | 0.92 | 6.5E-07 |
| H | 8.7 | < | 10.0 | 1.6 | 4.0 | 7.1E-06 | 1 | 3.6E-06 |
| I | 32.0 | | 32.0 | 0.4 | 0.0 | 1 | 1 | 0.89 |
| J | 10.0 | < | 15.0 | 5.2 | 5.0 | 0 | 1 | 1.7E-28 |
| J′ | 5.2 | < | 6.9 | 1.8 | 1.0 | 0.0014 | 1 | 0.00072 |
| K | 14.0 | > | 14.0 | -0.3 | -1.0 | 2.3E-14 | 1.2E-14 | 0.18 |
| K′ | 6.1 | | 6.1 | 0.0 | 0.0 | 1 | 0.96 | 0.95 |
| K″ | 3.8 | < | 4.8 | 0.9 | 0.0 | 0.0013 | 1 | 0.00067 |
| L | 19.0 | > | 18.0 | -0.6 | -1.0 | 5.6E-14 | 2.8E-14 | 0.95 |
| L′ | 3.1 | | 3.3 | 0.2 | 0.0 | 1 | 1 | 0.86 |
| β1-0 | 1.6 | > | 1.3 | -0.3 | -1.0 | 0.017 | 0.0087 | 0.93 |
| β1-1 | 4.5 | < | 5.8 | 1.3 | 2.0 | 1.6E-15 | 1 | 7.8E-16 |
| β1-2 | 4.5 | < | 5.8 | 1.4 | 2.0 | 1.9E-13 | 1 | 9.7E-14 |
| β1-3 | 4.3 | | 4.4 | 0.0 | 0.0 | 1 | 1 | 0.93 |
| β1-4 | 4.8 | > | 3.6 | -1.3 | -1.0 | 5.9E-06 | 2.9E-06 | 1 |
| β1-5 | 2.1 | > | 1.7 | -0.4 | 0.0 | 0.0031 | 0.0016 | 1 |
| β2-1 | 2.9 | | 2.9 | 0.0 | 0.0 | 0.89 | 0.72 | 0.51 |
| β2-2 | 2.9 | | 2.9 | 0.1 | 0.0 | 0.89 | 0.94 | 0.51 |
| β3-1 | 2.5 | | 2.6 | 0.1 | 0.0 | 0.0013 | 0.00066 | 0.0027 |
| β3-2 | 3.9 | < | 5.5 | 1.5 | 2.0 | 1.5E-13 | 0.95 | 7.4E-14 |
| β3-3 | 2.9 | < | 4.5 | 1.6 | 2.0 | 4.4E-16 | 0.78 | 2.5E-16 |
| β4-1 | 1.7 | | 2.2 | 0.5 | 1.0 | 0.062 | 1 | 0.031 |
| β4-2 | 2.0 | | 2.2 | 0.2 | 1.0 | 0.062 | 0.54 | 0.031 |
| β5-1 | 1.3 | | 1.1 | -0.2 | 0.0 | 0.87 | 0.49 | 1 |
| β5-2 | 1.3 | | 1.1 | -0.2 | 0.0 | 0.87 | 0.49 | 1 |
| β6-1 | 1.3 | | 2.0 | 0.7 | 1.0 | 0.43 | 0.98 | 0.22 |
| β6-2 | 1.1 | < | 3.2 | 2.2 | 3.0 | 0.032 | 1 | 0.016 |
